# Standard Issue: Copy number heterogeneity of JC virus standards discovered through next-generation sequencing

**DOI:** 10.1101/085415

**Authors:** Alexander L. Greninger, Allen C. Bateman, Ederlyn E. Atienza, Sharon Wendt, Keith R. Jerome, Linda Cook

## Abstract

Quantitative PCR is a diagnostic mainstay of clinical virology, and accurate quantitation of viral load among labs requires the use of international standards. However, the use of multiple passages of viral isolates to obtain sufficient material for international standards may result in genomic changes that complicate their use as quantitative standards. We performed next-generation sequencing to gain single-nucleotide resolution and relative copy number of JC virus (JCV) clinical standards. Strikingly, the WHO international standard and Exact^TM^ v1/v2 prototype standards for JCV showed 8-fold and 4-fold variation in genomic coverage between different loci in the viral genome, respectively, due to large deletions in the large T-antigen region. No such variation was seen for a clinical sample with high copy-number of JCV nor a plasmid control. Intriguingly, several of the JCV standards sequenced in this study with large T-antigen deletions were cultured in cell lines immortalized using SV40 T-antigen, suggesting the possibility of trans-complementation in cell culture. Using a cut-off of 2% variant allele fraction for junctional reads to define the presence of a strain, 11 different strains were present in the WHO standard. In summary, targeting of different regions of the same international standard could result in up to an 8-fold difference in quantitation. We recommend the use of next-generation sequencing to validate standards in clinical virology.

## Introduction

Polyomaviruses are unenveloped, double-stranded DNA viruses with a circular genome (1). Despite the discovery of multiple polyomaviruses over the last decade, only four polyomaviruses have been associated with human disease (1–5). JC virus (JCV) infection has been associated with progressive multifocal leukoencephalopathy (PML) in immunosuppressed patients (6, 7). JC virus has also been associated with two other neurological diseases, JC virus granule cell neuronopathy and JC virus encephalopathy (7).

Detection of JCV in the cerebrospinal fluid (CSF) is required for diagnosis of PML and quantitative levels of the virus have been associated with clinical outcomes of PML patients, with lower DNA values associated with better outcomes (8–11). However, viral loads in patients can vary dramatically, meaning analytically sensitive assays are required to protect against false negative tests (12). Real-time PCR is commonly used to measure levels of JCV in CSF (13).

Quantitative standards are of critical importance to real-time PCR in order to quantitate copies and compare different assays (14). The creation of laboratory standards in virology often relies upon growing up large amounts of virus in cell culture to recapitulate the viral particle in extraction and to ensure future preparations are not required. However, passaging a virus several times can create selection pressures that do not recapitulate viral biology in vivo. In this study, we used next-generation sequencing and confirmatory quantitative real-time PCR and droplet digital PCR to show that several standards used for JCV quantitative PCR contain multiple strains, some with large deletions in the T-antigen region, that incur large variability in measured copies depending on the locus measured. We hypothesize that these deletions are due to passage of the virus in SV40 T-antigen immortalized cell lines

## Materials and Methods

### JC virus moteriols

The 1st WHO International Standard for JC Virus DNA (NIBSC code: 14/114) was reconstituted in 1 mL of nuclease-free molecular-grade water and left for 20 minutes with occasional gentle agitation before use, according to the instructions. The instructions then state to dilute the international standard in the matrix routinely used within the laboratory for clinical diagnosis of JCV DNA, and that the diluted material should be extracted prior to JCV DNA measurement. We diluted the international standard 1:1 into 1ml of normal serum control (NSC; BioRad), followed by serial 10-fold dilutions in NSC prior to extraction.

The Exact Diagnostics (Fort Worth, TX) JCV prototype panel v1 and JCV prototype panel v2 each consist of six concentrations (1×10^7^, 1×10^6^, 1×10^5^, 1×10^4^, 1×10^3^, and 1×10^2^ copies/ml) of whole, intact JC virus MAD-4 strain. Each concentration was extracted and run on qPCR and ddPCR as described below.

JC virus ATCC strain 1397 was diluted for use as in-house JCV positive control. Control material was serial 10-fold diluted in NSC prior to extraction for ddPCR. For qPCR, the highest concentration was extracted and serial 10-fold dilutions were made in 10mM Tris-HCl.

A JCV-positive urine sample was serial 10-fold diluted in NSC prior to extraction and run on qPCR and ddPCR as described below.

Advanced Biotechnologies Inc (Eldersburg, MD) JC Human Polyomavirus (MAD1 strain) Viral qDNA PCR Control (17-943-500) was obtained at a stated concentration of 5×l0^6^copies/ml of DNA. This material was serial 10-fold diluted in 10mM Tris and run on qPCR and ddPCR without extraction as described below.

A plasmid containing the MAD-1 strain was ordered from Addgene (plasmid #25626), miniprepped, and diluted in TE buffer to approximately 10^6^ copies/ml (15). This material was then serial 10-fold diluted in 10mM Tris and run on qPCR and ddPCR without extraction as described below.

### DNA extraction and Quantitative Real-time PCR

The Roche MagNa Pure 96 (Roche, Indianapolis, IN) was used to extract DNA from the WHO, Exact, ATCC, and urine materials according to the manufacturers' instructions. The input volume was 500μl and the elution volume was 100μl. Extracted DNA was either used immediately or stored for up to 24 hours at 4°C before PCR.

The quantitative real-time PCR assay used was the clinical JCV assay currently run at the University of Washington Clinical Virology (16, 17). This assay targets the Large T region with a single primer pair and a FAM-TAMRA probe. WHO dilutions were named as standards used for converting quantification cycle (Cq) to copies/ml.

Real-time quantification was also performed using the Focus Diagnostics master mix from DiaSorin (Cypress, CA) which utilizes JCV primer pair (MOL9021) and 2.5X Universal Master Mix. The primer pair includes a Scorpion primer/probe and targets a conserved region of VP2/3 gene, as described below. WHO dilutions were used to convert quantification cycle (Cq) to copies/ml in post amplification analysis.

### Droplet digital PCR

Prior to ddPCR, restriction enzyme digestion was performed with HindIII (New England Biolabs, Ipswitch, NY). To 5μl of extracted DNA, 3μl of water, 1μl of 10× CutSmart buffer, and 1μl of HindIII (20,000U/ml) were added, followed by incubation at 37°C for 1 hour. After incubation the mixture was diluted 1:5 by the addition of 40μl of water, and 10μl of this dilution was used per PCR reaction.

Droplet digital PCR was performed using the BioRad system. BioRad ddPCR mastermix, primers, and probes (final primer and probe concentrations equal to final qPCR concentrations) were added to a 96-well plate and vortexed, followed by droplet generation on the QX100 droplet generator. Droplets were transferred to a 96-well PCR plate and amplified on a 2720 Thermal Cycler (Applied Biosystems) with the following thermocycling parameters: 94°C for 10 min, followed by 40 cycles of 94°C for 30 sec and 60°C for 1 min, and a 98°C hold for 10 min. Droplets were read in the QX200 droplet reader (BioRad) immediately following amplification. Data were analyzed with QuantaSoft analysis version 1.3.2.0, and quantification was calculated to reflect copies/ml of the initial specimen.

### Next-Generation Sequencing

Because of the low concentration of the JCV standards (0.1-0.2 ng/uL), DNA from the standards was used neat for dual-indexed Nextera XT sequencing library preparation followed by 18 cycles of amplification (18). Strains from WHO, Exact^TM^ v1, pMAD1 plasmid, as well as a clinical strain from urine were sequenced on a single-end 185bp run on an Illumina MiSeq. JCV-1397 from ATCC, WHO, Exact^TM^ v1, Exact^TM^ v2, and the urine clinical strain were also sequenced on a 2×260bp run on an Illumina MiSeq. Sequencing reads were adapter-and quality-filtered (Q30) using cutadapt and aligned to the JC virus reference genome (NC_001699) using Geneious v9.1.4 mapper with structural variant detection enabled (19–21). Coverage maps were produced from bam files generated using Geneious using the genomecov option from bedtools (22).

To determine the locus used in the Focus Diagnostics analyte specific-reagent, an amplicon created via real-time PCR with the Focus primers was subjected to half-reaction of end-repair and dA-tailing, followed by adapter ligation and PCR amplification with Truseq adapters, using a Kapa HyperPlus kit. The indexed amplicon was sequenced on a 2×500bp MiSeq run and reads were adapter/quality-trimmed and mapped to the JC virus reference genome as above. The location of the primers is depicted in Figure 1B.

**Figure 1 –.**
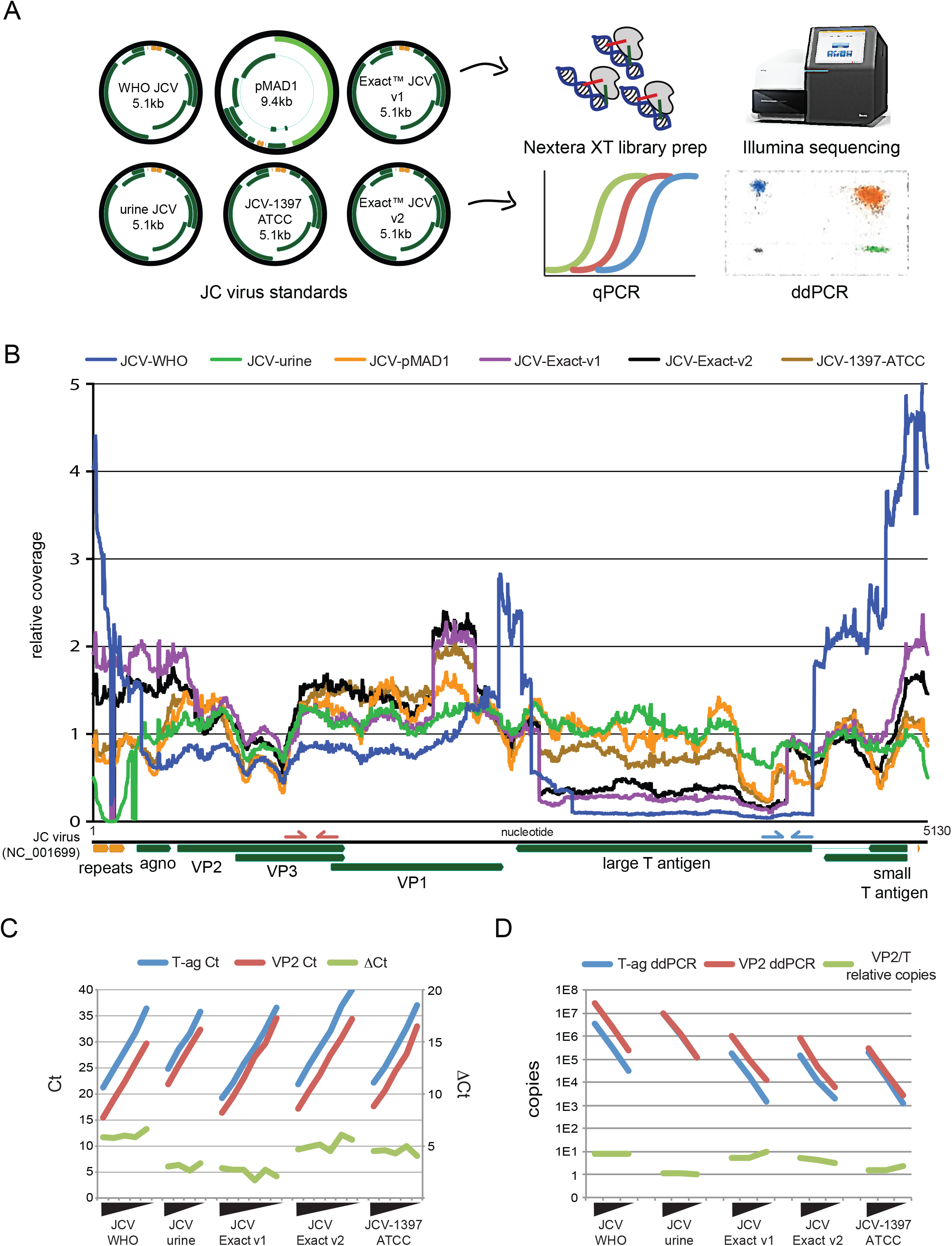
Next-Generation Sequencing of JC Virus Standards Reveals Deletions and Copy Number Heterogeneity. A) Six different JC virus materials were deep sequenced and five standards were tested by qPCR and ddPCR. Gene organization of each JC virus is shown in green, with regulatory regions depicted in orange. Of note, the 9.4kb pMAD1 plasmid inserted the backbone at the location of the probe used in the T-antigen qPCR and ddPCR assay and cannot be quantitated at that locus. B) Coverage plot of six different standards of JC virus mapped to the JC virus NCBI reference genome (NC_001699). Three of the six standards include a large deletion in the T-antigen region that constitutes a greater than 4-fold difference in copy number relative to the structural genes. Reductions in coverage in the regulatory repeat region are due both to small deletions and sequence divergence relative to JC virus reference genome. Primers for the Focus PCR analyte-specific reagent targeting the VP2/3 region are shown on the JC virus genome in red, while the pep primers targeting the T-antigen region are shown in blue. C and D) Confirmation of the copy number differences seen by sequencing was performed with qPCR (C) and ddPCR (D) using PCR primers against the VP2/3 gene and T-ag gene.

## Results

Based on our previous discovery of copy number heterogeneity in the BK virus international standard, we performed next-generation sequencing on a variety of JCV standards present in our laboratory. We obtained a provisional WHO standard for JCV, two different versions of the Exact^TM^ (v1 and v2), a urine specimen with a high copy number of JCV present, the ATCC JC virus strain 1397, and the original 1980 plasmid with the Mad-1 strain of JC virus cloned into it. All strains were sequenced using Nextera XT libraries on an Illumina MiSeq. qPCR and ddPCR using the Focus PCR primers targeting the VP2/3 region and the UW Virology clinical primers targeting the T-antigen region were performed as well (Figure 1A) (ref for primers).

Deep sequencing of the JC virus standards resulted in between 0.4-80% of the sequencing reads mapping to the JC virus (Table 1). Sequencing of DNA extracted from a urine specimen that was known to contain a high copy number of JC virus, as well as DNA extracted from a plasmid containing cloned JC virus both yielded the lowest variation in coverage (Figure 1B, Table 1). qPCR and ddPCR quantitation of the clinical urine specimen indicated equal copy numbers at the VP2/3 and T-antigen loci of these materials.

**Table 1 –.** Next-Generation Sequencing Results of JC Virus Standards

| **Strain** | **Total Reads** | **JC virus reads** | **Mean Coverage** | **CV** | **Number of deletion variant strains present (highest variant frequency)¶** |
| --- | --- | --- | --- | --- | --- |
| WHO | 512,609 | 406,100 | 13445 | 98.4% | 11 (31%) |
| JCV urine | 299,886 | 59,115 | 1574 | 26.0% | 0 |
| pMAD1 plasmid | 2,460,512 | 963,917 | 26060 | 32.5% | 0 |
| Exact™ v1 | 586,222 | 96,765 | 2943 | 57.8% | 6 (34%) |
| Exact™ v2 | 2,602,925 | 9,860 | 383 | 54.3% | 2 (27%) |
| ATCC-1397 | 1,694,258 | 11,527 | 393 | 39.1% | 3 (18%) |

¶ Deletion variant strain defined by deletion >50bp present at 2% variant frequency

The WHO international standard contained an approximate 8-fold variation in coverage between structural genes and the much of the T-antigen region and contained the largest coefficient of variation in coverage among the strains at 98.4% (Figure 1B and D). Analysis of junctional reads indicating deletions of >50bp present at greater than 2% variant allele frequency revealed 11 different strains present in the WHO international standard. The Exact^TM^ v1 and v2 standards and the ATCC-1397 standard each contained multiple strains, including a variant strain present at 18-34% abundance that contained the same duplication of the C-terminus of the VP2/3 gene inserted into a deletion C-terminus of the T-antigen (Table 1). Of note, both of the Exact^TM^ strains as well as the ATCC-1397 strain were grown in the Cos-7 cell line. The Exact^TM^ v1 and v2 standards both yielded an approximate 4-to 5-fold difference in copy number between structural genes and T-antigen, while the ATCC-1397 strain had a 1.5-fold difference in copy number between these loci (Figure 1C/D).

## Discussion

Using deep sequencing and confirmatory qPCR and ddPCR, we show four different qPCR standards for JC virus contain strains with large, different deletions in the T-antigen region. These deletions result in significant copy number differences between different loci in the JC virus genome that could lead to radically different sensitivities and quantities being reported by various JC virus qPCR assays. By contrast, deep sequencing of a clinical specimen with JC virus and of a plasmid containing JC virus sequence do not show the large differences in copy number found in the cell-culture adapted strains. These results mirror those we recently reported in an international standard for BK virus (Bateman et al. (2016), under review).

Of note, one of the JC virus standards sequenced in this study was cultured in the Cos cell line. The Cos cell line is derived from a primary monkey cell line that was transformed with SV40 T-antigen and expresses both large T antigen and small t antigen (23). Given that the deletions seen in this study occurred in the T-antigen region of the JC virus, were seen in approximately 90% of the JC virus strains present, and that T-antigen is known to be required for polyomavirus replication, we hypothesize that the SV40 T-antigen in the Cos cell line is providing nonstructural functions such as DNA binding, unwinding, viral replication, and transformation in trans to the JC virus strains with T-antigen deletions sequenced in this study. Indeed, SV40 T-antigen has been shown to be required for archetype JC virus replication, as JC virus is unable to replicate in the untransformed Cos parental cell line CV-1 (24). Thus, Cos cells may have been specifically chosen for their ability to support high levels of JC virus growth given the need for large amounts of highly concentrated virus to provide qPCR standards to multiple labs around the world, albeit with the unforeseen consequence that the cell line would yield JC virus strains with large deletions.

The large coverage differences in the T-antigen portion of the JC virus materials led us to investigate knock-on effects due to the function of the large T antigen in polyomavirus replication. The many functions of the large T-antigen include binding multiple host proteins (including p68 DNA polymerase/primase, p53, Rb, hsc70, and replication protein A), binding to itself to form hexamer, binding the origin of replication, unwinding DNA, and translocating to the nucleus (25, 26). Of these, only origin binding would be potentially compromised by complementation. BK, JC, and SV40 viruses have different nucleotides flanking the pentanucleotide GAGGC motif used for double hexamer formation at the origin that are not involved in hairpin formation (SV40: gaggcCgaggc, JCV: gaggcGgaggc, BKV: gaggcAgaggc) (1). Interestingly, the BKV sequence recovered in Bateman et al. (2016) showed a 5:1 variant allele frequency at the flanking nucleotide for the SV40 origin sequence compared to the BKV origin sequence. This variant allele frequency matched the copy number difference found in the BKV large T-antigen. No sequence alteration was found in the origin for either of the JC virus sequences isolated in this study, although a tatataGaaaa change (from tatataTaaaa) was found in the downstream TA-rich region that to date has only been recovered in one JC virus ever deposited in NCBI (KF788289) (27). This data would be consistent with the hypothesis that SV40 large T-antigen shows preference for its own origin relative to BKV but not JCV, as the BKV origin changed to the SV40 origin but the JCV origin did not change.

The recovery of JC and BK virus sequences with large deletions in cell culture is likely due to the unique biology and sequence identity of the polyomaviruses. Encapsidation of viral DNA is thought to be dependent on viral structural proteins with no contribution from viral non-structural proteins beyond DNA replication (28). The amino acid sequence conservation between SV40 and JC and BK virus T-antigen is approximately 73%. As described above, the origin DNA sequence is nearly identical between different polyomaviruses with the nucleotide sequence required for T-antigen binding being absolutely conserved. JC virus T-antigen J domain can replace SV40 J-domain and retain replication activity (29). Chimeras of JC virus and SV40 showed that viruses containing JC virus regulatory sequences and SV40 coding regions could replicate, consistent with the JC virus recovered in this study (30). Thus, polyomavirus non-structural proteins show the ability to complement each other (31, 32). Indeed, different deletions were recovered in three of the cell culture-associated BK and JC polyomavirus sequenced.

Our study shows the importance of deep sequencing of standards to validate reagent integrity before they are scaled internationally (33). Deep sequencing provides single-nucleotide resolution of strains present and the relative copy number of loci across the genome. Deep sequencing of these standards demonstrated the presence of multiple strains present with radically different copy number due to deletions, as well as single-nucleotide changes that may affect PCR primer binding and overall quantitation, making growth of BK and JC virus in SV-40 transformed cell lines a potentially suboptimal choice for a qPCR international standard.

Supplemental Table – **qPCR and ddPCR data table depicted in Figure 1C and 1D**.

